# CESAR: A R Package for High-Sensitivity Detection of Copy Number Variations in ctDNA Using Segmentation and Anchor Recalibration

**DOI:** 10.64898/2026.03.09.710442

**Authors:** Jinchang Lu, Shuai Ni, Kui Lu, Bo Wu, Yun Wang, Lina Wang, Ning Wu, Xiaobing Jiang

## Abstract

**Background:** Detecting copy number variations (CNVs) in circulating tumor DNA (ctDNA) is crucial for the companion diagnosis and resistance monitoring of various solid tumors (e.g., NSCLC, Glioblastoma). However, when tumor-derived DNA fractions are extremely low (often <1%), traditional depth-based methods frequently fail due to non-linear sequencing depth fluctuations and probe-specific capture biases inherent to targeted Next-Generation Sequencing (NGS).

**Methods:** We developed CESAR (CNV Estimation with Segmentation and Anchor Recalibration), a computational tool for tumor-only CNV detection in targeted NGS panels. CESAR uses Circular Binary Segmentation (CBS) to re-partition target regions according to relative capture efficiency, then applies a data-driven “anchor” selection procedure that, for each target segment, identifies a personalized set of co-varying genomic segments. By selecting the anchor set that minimizes the coefficient of variation (CV) of the anchor-recalibrated depth ratio across a panel of normals, CESAR recalibrates the per-segment baseline and suppresses probe-specific technical noise. Copy-number status is then called from the deviation of the observed ratio against this trained baseline.

**Results:** Using standard DNA reference materials, CESAR identified amplifications of MET, ERBB2, and EGFR at low tumor fractions. CESAR resolved focal alterations as subtle as 2.18 copies (a 1.09-fold change relative to the diploid baseline) while reporting no false-positive amplifications in diploid control regions. Applied to a clinical cohort of nine NSCLC ctDNA samples profiled on a 94-gene panel, CESAR successfully separated three MET-amplified samples from six matched negatives, where the standard CNVkit-based pipeline failed to distinguish them. In head-to-head benchmarking on identical data, CESAR reduced technical variance relative to the widely used CNVkit and more reproducibly resolved low-level copy-number gains, particularly in the depth-heterogeneous MET amplicon.

**Conclusions:** CESAR provides a stable and sensitive framework for tumor-only CNV calling in liquid biopsies. On reference standards it outperforms CNVkit in bias and reproducibility, and on a clinical NSCLC cohort it recovered MET CNV abnormalities that a standard CNVkit pipeline missed. Validation on larger and more diverse cohorts is warranted.

## 1. Introduction

Genomic instability is a fundamental hallmark of cancer, frequently manifesting as Copy Number Variations (CNVs)—the amplification or deletion of chromosomal segments ranging from kilobases to entire chromosome arms [1]. Across human malignancies, somatic CNVs serve as primary drivers of oncogenesis by either hyperactivating oncogenes through focal amplifications or inactivating tumor suppressor genes via deep deletions [2]. In numerous malignancies, specific focal CNVs are not merely incidental passenger mutations but critical, actionable clinical targets. For instance, in Non-Small Cell Lung Cancer (NSCLC), the amplification of EGFR and ERBB2 initiates oncogenic cascades, while acquired MET amplification serves as a predominant bypass resistance mechanism against targeted therapies [3, 4]. Focal CNVs are similarly central to other solid tumors: in diffuse gliomas, alterations such as EGFR amplification and the homozygous deletion of CDKN2A/B inform both the current World Health Organization (WHO) diagnostic classification and treatment eligibility [5, 6]. Accurate, real-time monitoring of such CNVs via liquid biopsy is therefore valuable across diverse solid tumors for dynamic therapeutic decision-making [7, 21].

Traditionally, detecting tumor genomic alterations relied on invasive tissue biopsies. In recent years, liquid biopsy using circulating tumor DNA (ctDNA)—fragmented, tumor-derived DNA circulating in peripheral blood or cerebrospinal fluid (CSF)—has substantially advanced molecular oncology by offering a minimally invasive, real-time snapshot of the tumor genome [8]. However, analyzing CNVs from ctDNA presents a substantial statistical and technical challenge. The primary obstacle stems from the inherently low abundance of tumor-derived DNA in the bloodstream. In the vast majority of early-to-mid stage advanced cancer cases, ctDNA constitutes less than 10%—and frequently falls below 1%—of the total cell-free DNA (cfDNA) [10].

To detect such ultra-low fraction CNVs, whole-genome sequencing (WGS) is mathematically and economically unfeasible at extreme depths. While low-coverage WGS combined with statistical segmentation models can accurately detect megabase (Mbp)-level large-scale CNVs [20], this approach is generally limited to patients where the ctDNA fraction exceeds 10% [7, 8]. Consequently, targeted CNV detection using highly focused NGS panels has become the clinical standard. By restricting the sequencing footprint to a targeted subset of clinically relevant genes (e.g., a 30 kb panel), sequencing capacity can be concentrated to achieve ultra-deep coverage (often exceeding 5,000x to 10,000x) [9]. This immense depth theoretically allows for the statistical quantification of minute increases in reads caused by focal gene amplifications in ctDNA.

Despite the utilization of ultra-deep targeted panels, robust CNV detection in liquid biopsy remains difficult due to the intrinsic, non-biological depth fluctuations and technical noise inherent to the NGS capture and library preparation processes [10]. Current computational methodologies for tumor-only CNV calling predominantly rely on three strategies, each exhibiting significant limitations when applied to highly restricted capture regions [17]. The most common approach compares the sequencing depth of a tumor sample against a Panel of Normals (PoN) [11, 12, 16]. Yet, this method assumes a linear correlation between individual probe coverage and overall sample depth, a premise shown to be unreliable given probe-specific capture biases [13]. An alternative method utilizes the B-allele frequency (BAF) distribution of heterozygous single nucleotide polymorphisms (SNPs). While sensitive when the target region contains on the order of 50 heterozygous SNPs [14], BAF-based detection is statistically underpowered for small, focused panels that do not encompass enough SNPs. Lastly, directly detecting the chimeric sequences formed at the junction points of structural rearrangements (typically spanning 1 to 100 Mbp) offers high single-molecule sensitivity in principle [15]. However, the probability of capturing these rare junction reads within the limited footprint of a 30 kb targeted panel is very small.

To address these limitations, we developed CESAR, a method designed for sensitive, tumor-only CNV detection in targeted ctDNA panels. Rather than relying on global average-depth normalization, CESAR re-partitions the targeted regions based on relative capture efficiency and identifies a personalized set of “anchor” segments for each target CNV gene. By modeling the ratio between the target depth and its matched anchors, CESAR mitigates systematic, probe-specific variance, enabling stable and statistically supported detection of gene-level CNVs (e.g., MET, ERBB2, EGFR) even when the ctDNA fraction is low.

---

## 2. Materials and Methods

The CESAR algorithmic framework is explicitly designed to fulfill two primary computational objectives: (1) Capturing intra-sample variance: re-partitioning the targeted panel (e.g., 30 kb) based on probe location and capture efficiency to precisely measure regional probe depth; and (2) Reducing inter-sample variance: substituting the global average depth with dynamically matched “anchor” regions when calculating relative depth, thereby improving the stability of CNV measurements across different samples.

### 2.1. Re-segmentation Based on Relative Capture Efficiency

In targeted sequencing, the fundamental assumption that sequencing depth increases linearly with total sample coverage is often violated due to probe-specific capture biases. To capture this non-linear variation, CESAR initiates a learning phase utilizing a cohort of normal control samples (Panel of Normals, PoN) that lack copy number variations. Using a PoN is a well-established strategy to establish baseline capture efficiencies and filter recurrent sequencing artifacts [11, 12].

Based on the average sequencing depths across all normal controls, the known targeted regions are initially divided into micro-segments of approximately 40 base pairs (bp). We employed the Circular Binary Segmentation (CBS) algorithm [18] to identify change points in the normalized read depth profile. By evaluating the depth similarity and segment size of adjacent micro-segments, contiguous regions with congruent capture efficiencies are merged. Post-merging, the entire targeted panel is re-partitioned into final segments ranging from 100 to 400 bp. These empirically derived segments serve as the foundational genomic units for downstream anchor identification. By executing this step, CESAR effectively captures and delineates the inherent capture-efficiency discrepancies among different probe regions within a single sample. Furthermore, to maximize analytical flexibility, the software allows users to bypass this automated learning phase by providing a custom, user-defined BED file for subsequent downstream operations.

### 2.2. Dynamic Anchor Recalibration Algorithm

The central step of CESAR mitigates the systematic errors induced by referencing global average depth. We observed that when the total sample depth doubles, the depth of certain probe regions increases by more than twofold. Consequently, using the global average depth as a universal reference for single-probe coverage changes introduces systematic errors.

To address this, CESAR draws on count-data models that favor dynamically optimized reference sets over global averages (e.g., ExomeDepth [19]). However, rather than selecting a reference cohort at the sample level, CESAR applies a localized approach, automatically identifying a tailored set of intra-panel “anchor” segments for each target CNV gene (e.g., MET, ERBB2, EGFR). For a given target segment T, CESAR evaluates all other segments within the panel across the normal training cohort to find a subset of N segments whose depth variation trend most closely mirrors that of segment T.

Mathematically, CESAR first computes the Pearson correlation matrix for the sequencing depths of all segments across the training samples. For the target segment T, all candidate anchor segments are ranked in descending order based on their correlation coefficients. CESAR then iteratively evaluates candidate anchor sets A(T,N) = {S1, S2, …, SN} by varying the number of top-ranked segments (N). The algorithm automatically searches for the number of segments whose variation trend most closely matches the target, so as to minimize inter-sample variance. Selecting too few anchors makes the baseline sensitive to noise in individual segments, while including weakly correlated segments reintroduces the very variance the anchors are meant to suppress. CESAR therefore bounds N within a user-configurable range and selects the value that minimizes the coefficient of variation (below), guarding against both extremes. Same-gene segments are excluded from the candidate pool by default, so that a whole-gene amplification cannot recruit its own amplified segments as anchors.

Let $R_{T,j}$ be the ratio of the mean depth of the candidate anchor set to the target segment’s depth in the $j$-th sample:

R(T,j) = [Average of Depth(S(k,j)) for k=1 to N] / Depth(T(j))

For each subset size N, CESAR calculates the coefficient of variation (CV = sigma/mu) of R(T) across all training samples and selects the number of anchors N that minimizes it. Substituting the target region’s global expected depth with this custom, highly correlated anchor depth improves the stability of CNV measurements and limits systematic background variance. Defining the ratio this way—anchors over target—means that a copy-number gain in the target lowers R(T), and the reported copy number ($\mu / R_{obs}$, Section 2.3) rises above the diploid baseline accordingly.

### 2.3. Statistical Modeling and CNV Calling

Following the identification of optimal anchors, CESAR establishes a baseline statistical model for the relative depth ratios (R(T)) derived from the normal cohort. For every target segment, we fit a Normal distribution N(mu, sigma^2) to the ratios observed across the training samples, estimating mu and sigma by maximum likelihood.

During the detection phase, for a given test sample (e.g., a patient’s ctDNA), CESAR computes the observed relative depth ratio r(obs) between the target region and its learned anchor regions. This value is converted to a Z-score against the trained baseline, z = (r(obs) - mu)/sigma, and a two-sided P-value is obtained as P = 2Phi(-|z|). To avoid numerical underflow at extreme significance, P*-values are floored at a small constant (default 10^(-100)) before the -log10 transformation used to report a confidence score. The estimated relative copy number is reported as mu/r(obs), so that a value of 1 corresponds to no change from the diploid baseline. Because the test operates on the anchor-recalibrated ratio rather than raw depth, it is robust to the probe-specific, non-linear depth fluctuations described above.

Per-segment calls are aggregated to the gene level by averaging the estimated copy ratios and confidence scores across all segments annotated to a given gene. CESAR currently reports pergene significance without a formal multiple-testing correction across genes; for panels interrogating many independent loci, users should interpret confidence scores accordingly.

---

## 3. Results

### 3.1. Re-segmentation of the Target Capture Space

To address the inherent heterogeneity of hybridization probes in target enrichment sequencing, we first applied the Circular Binary Segmentation (CBS) algorithm to re-partition the theoretical target regions. Unlike relying on arbitrary genomic coordinates, this approach dynamically grouped probes based on their empirical relative capture efficiencies across the Panel of Normals (PoN). As illustrated in Figure 1, probes with differing capture efficiencies were successfully demarcated into distinct analytical segments. This empirically driven segmentation allowed for a more precise evaluation of efficiency discrepancies among diverse probes within the same sample, laying the structural foundation for downstream recalibration.

**Figure 1.**
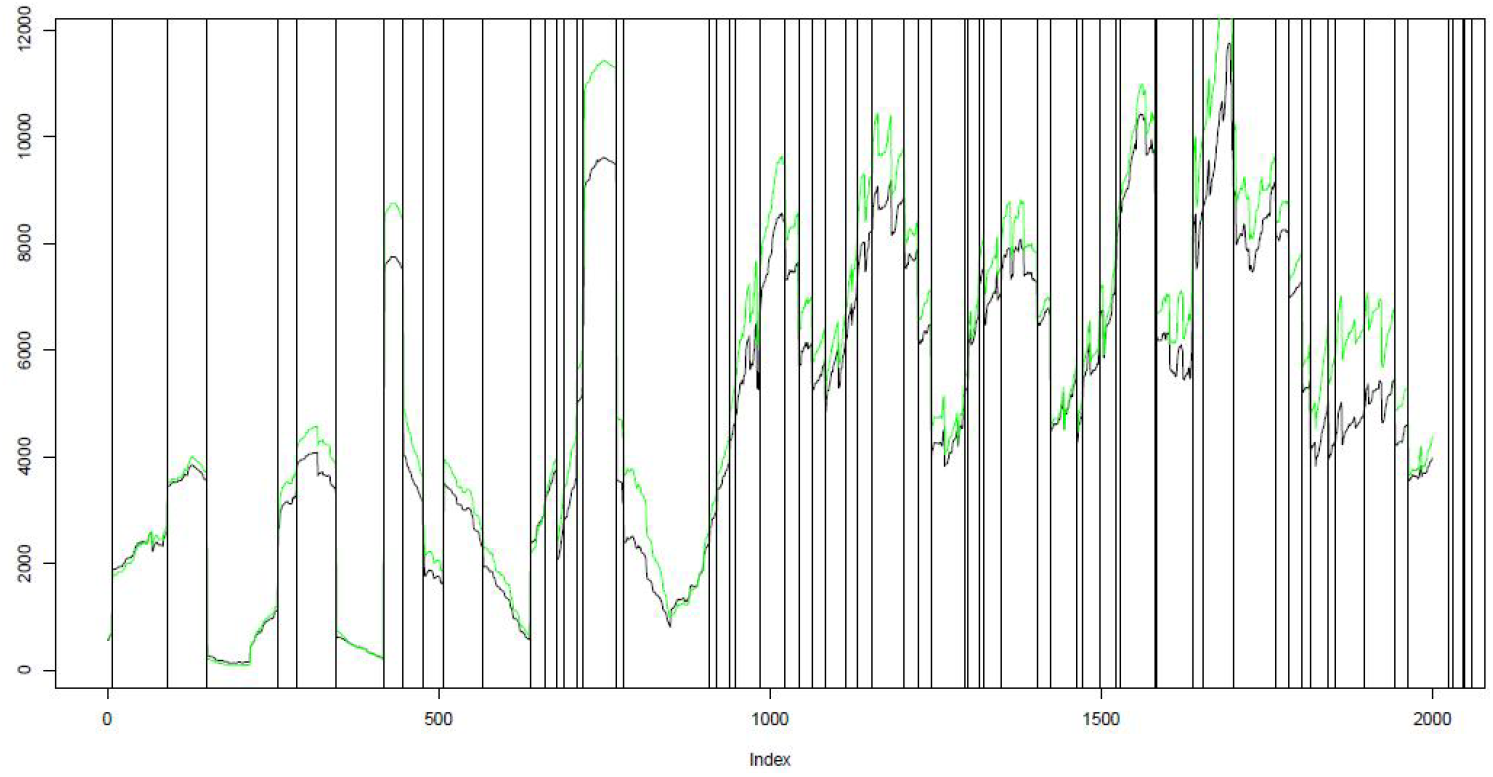
CBS-based re-segmentation of the target capture space. (Insert segmentation result plot here. Legend: The x-axis represents the genomic coordinates covered by the hybridization probes, and the y-axis represents the corresponding sequencing coverage depth. The plot visualizes the CBS algorithm successfully partitioning a ∼2,000 bp contiguous target region into approximately 60 distinct analytical segments (yielding an average segment length of ∼33.3 bp). This high-resolution segmentation effectively distinguishes sub-regions with drastically different sequencing depths. These sharp inter-segment depth discrepancies are inherently driven by probe-specific hybridization kinetics and local sequence contexts, representing biological and physical artifacts that are notoriously difficult to eliminate during library preparation. By delineating these boundaries, CESAR successfully isolates and characterizes probe-level variance.)

### 3.2. Reduction of Systematic Variance via Anchor Recalibration

Following segmentation, we addressed the non-linear relationship between individual probe coverage and total sample depth. By substituting the global average depth with the dynamically selected “anchor” depth for relative CNV calculation, CESAR improved measurement stability across samples. As depicted in Figure 2, replacing the average depth with the anchor depth reduced the variance of the MET gene’s relative depth ratio across samples by more than 50%. This stabilization separates true CNV signals from the background baseline, which is especially important when tumor fractions are low.

**Figure 2.**
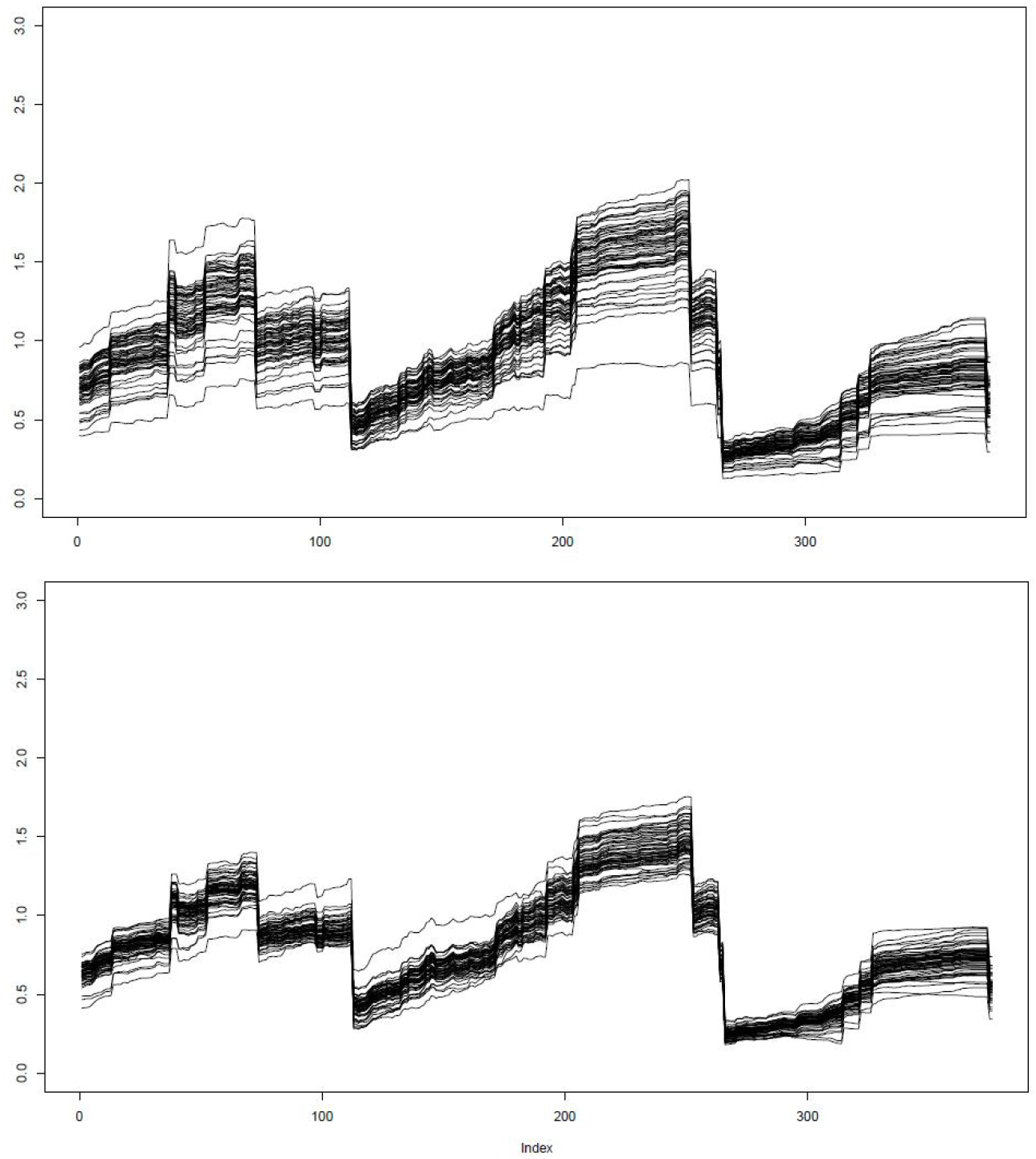
Comparison of relative depth ratio variance. (Insert variance comparison plot here. Legend: The upper panel demonstrates the non-linear correlation between specific regional coverage and global average coverage. The lower panel compares the fluctuation of the MET gene relative depth ratio before and after anchor recalibration, showcasing a >50% reduction in inter-sample variance.)

**Figure 3.**
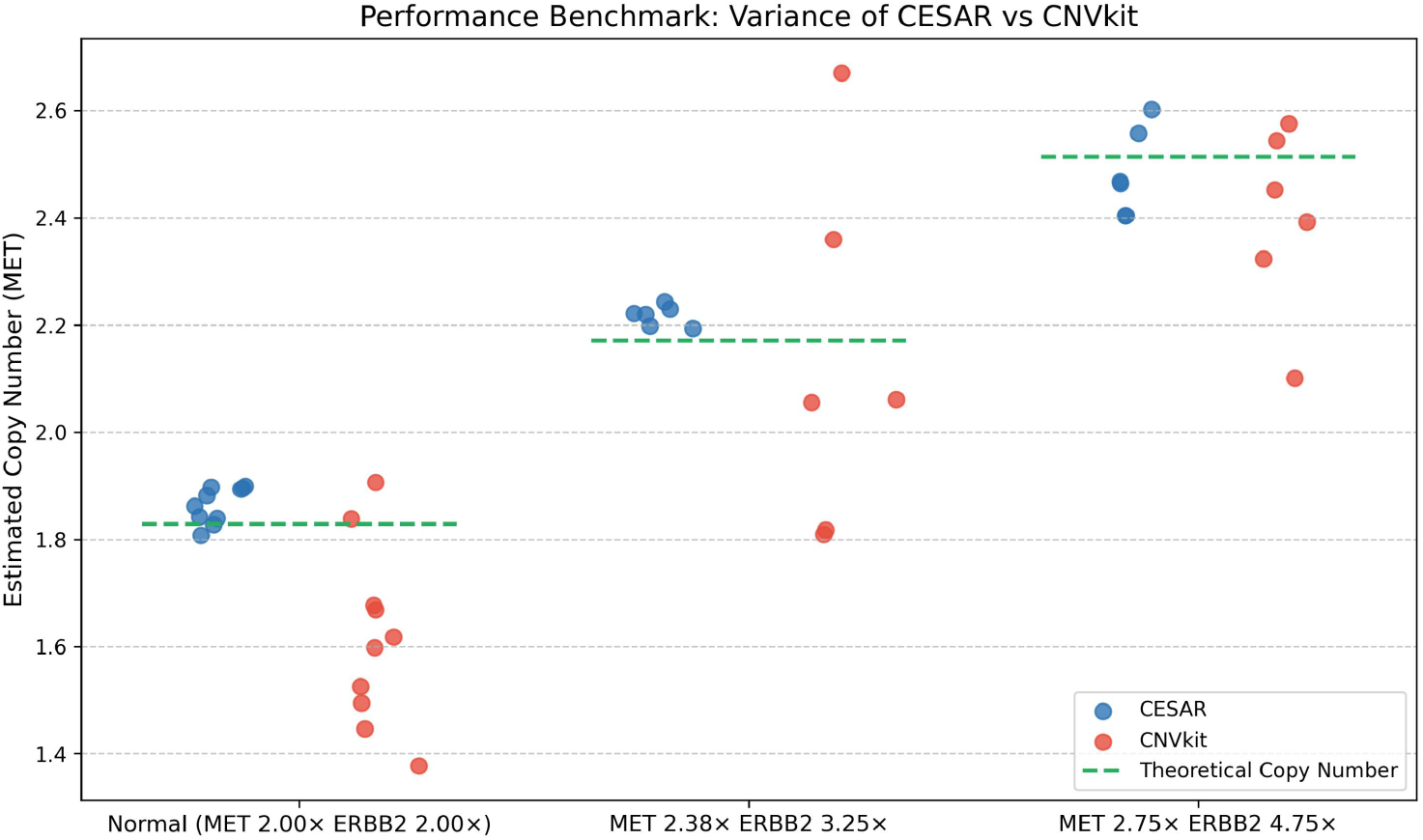
Performance comparison between CESAR and CNVkit on the MET gene. (Insert benchmark plot here. Legend: The plot contrasts the detection variance between CESAR and CNVkit for the highly heterogeneous MET amplicon. CESAR shows tighter clustering around the theoretical copy numbers than CNVkit, whose estimates spread more widely.)

### 3.3. Performance Validation and Benchmarking Using DNA Standards

To rigorously evaluate the stability and sensitivity of CESAR, we designed four distinct batches of standard reference materials (Table 1) representing a severe challenge for conventional callers.

**Table 1.** Experimental design of DNA standard reference materials.

| Batch | Sample Size (n) | MET Copy Number | ERBB2 Copy Number | EGFR Copy Number |
| --- | --- | --- | --- | --- |
| Batch 1 | 10 | 2.00× (Diploid) | 2.00× (Diploid) | 2.00× (Diploid) |
| Batch 2 | 6 | 2.18× (Amplification) | 2.625× (Amplification) | 2.00× (Diploid) |
| Batch 3 | 6 | 2.375× (Amplification) | 3.25× (Amplification) | 2.00× (Diploid) |
| Batch 4 | 6 | 2.75× (Amplification) | 4.75× (Amplification) | 2.00× (Diploid) |

Using the diploid samples of Batch 1 as the panel of normals to train the anchor sets and baseline ratio distributions, CESAR identified all of the true CNV alterations across the three amplification testing batches (Batches 2–4) and estimated copy numbers close to the engineered values. In the negative control regions (EGFR, diploid across all batches), CESAR reported no false-positive amplifications, indicating stable behavior on true-negative targets.

We subsequently benchmarked CESAR against the widely used depth-based tool CNVkit on the identical dataset. As detailed in Tables 2–4, both tools performed comparably in detecting high-level amplifications (e.g., ERBB2 ≥ 4.75x; Figure 4), with sensitivity and specificity summarized in Table 3. However, CNVkit showed clear limitations in identifying low-level copy-number gains, particularly for the MET gene.

**Figure 4.**
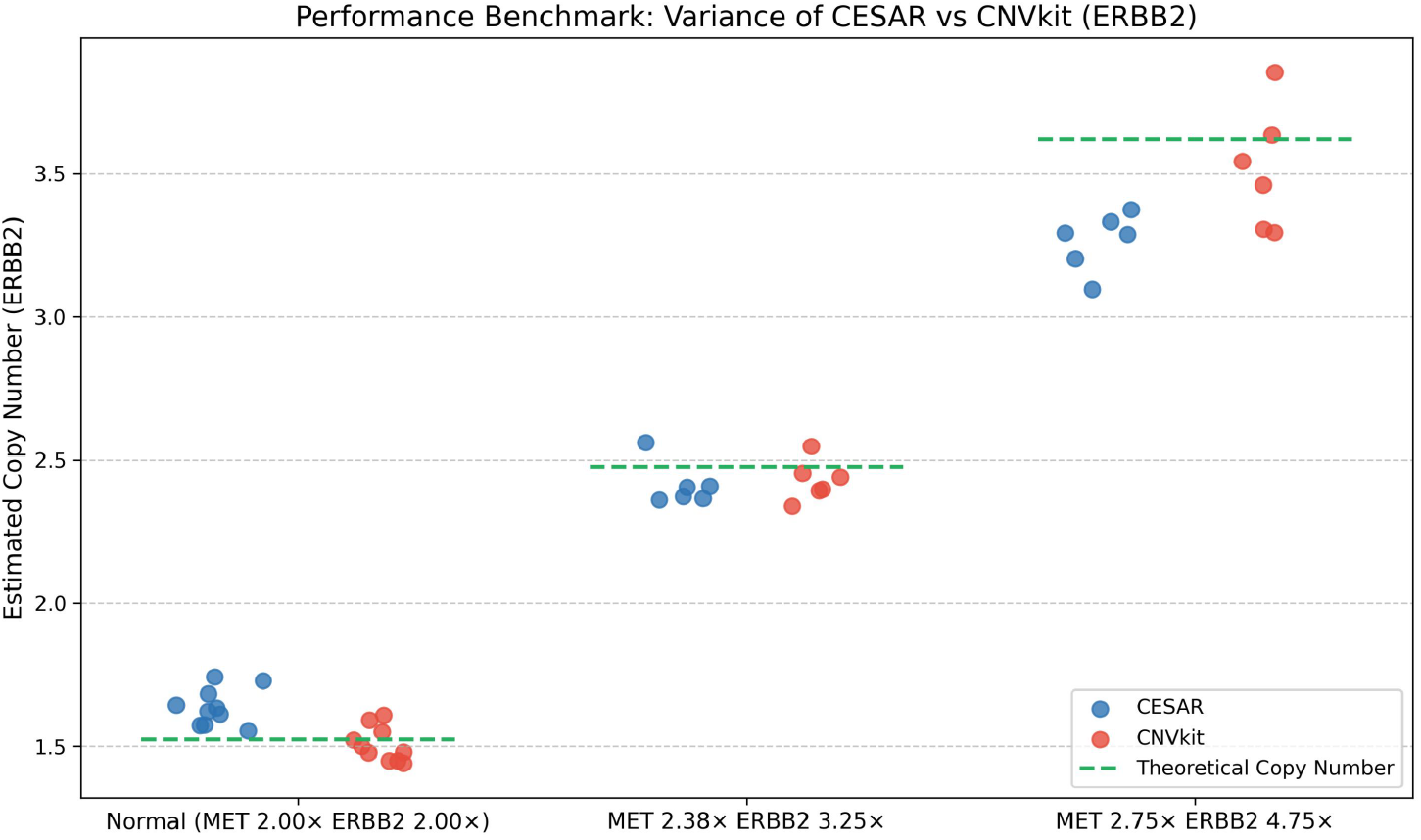
Performance comparison between CESAR and CNVkit on the ERBB2 gene. (Insert ERBB2 plot here. Legend: While both tools successfully identify high-level ERBB2 amplifications, CESAR maintains a noticeably tighter variance and less deviation from the theoretical copy number baseline compared to CNVkit.)

In the normal control group (true MET copy number = 2.0), CNVkit underestimated the copy number by an average of 0.21 copies (SD = 0.17), whereas CESAR’s bias was near zero (+0.04, SD = 0.03). In the moderate amplification cohort (Batch 3, MET 2.375x, n = 6), CNVkit’s inter-sample variance was roughly 17 times higher than CESAR’s (SD = 0.33 vs. 0.02), indicating substantially lower reproducibility. Within this same MET 2.375x group, CNVkit’s estimates ranged from 1.818 to 2.670—a maximum intra-group spread of 0.852 copies, which spans the standard clinical threshold for amplification. For the same 6 samples, CESAR’s estimates fell between 2.194 and 2.244, a maximum spread of 0.050 copies (Figure 3). This indicates that CNVkit’s reproducibility across identical standards is insufficient for the tight thresholds used in clinical ctDNA applications, whereas CESAR’s is not.

The inferior performance of CNVkit on MET is intricately linked to the sequencing depth heterogeneity within the amplicon. The MET target region exhibited a massive 3.64-fold depth fluctuation across loci (1,619x–5,900x), significantly higher than the 2.50-fold fluctuation in the ERBB2 region (3,508x–8,782x). CNVkit relies on per-locus depth normalization and subsequent CBS segmentation; when an amplicon exhibits such steep internal depth gradients, this normalization becomes highly unstable, introducing technical noise that concurrently inflates both bias and inter-sample variance. Furthermore, CNVkit lacks an inherent statistical significance testing framework, struggling to differentiate authentic low-level gains from noise. In contrast, CESAR does not require absolute depth uniformity: it models each target segment’s depth ratio against a set of co-varying anchor segments and evaluates deviations with a Z-score against the trained baseline distribution. Because the anchors track the same non-linear coverage behavior as the target, this recalibration reduces both bias and inter-sample variance relative to CNVkit in these heterogeneous targeted-sequencing data.

Table 2. Overall CNV detection results across all samples by CESAR and CNVkit.

Table 3. Detection sensitivity and specificity of CESAR versus CNVkit.

Table 4. Intra-group CNV detection stability and variance for identical reference standards.

### 3.4. Illustration of the Anchor Selection Process

To illustrate the anchor recalibration step, we visualized the anchor selection process for a representative MET segment (Figure 5). The top-left panel shows the distribution of Pearson correlation coefficients between the target segment and all other segments in the panel; the selected anchor segments (marked in red) exhibit the highest correlations (>0.9), indicating that they track the target’s depth variation most closely. The top-right panel shows the CV minimization criterion: as the number of anchors increases from 5 to 30, the coefficient of variation decreases and then plateaus, and CESAR selects the count that minimizes the CV (13 anchors in this example). The bottom panel confirms the stability of the anchor-recalibrated depth ratio across the six training samples: all ratios cluster tightly around the mean (6.09) with a CV of 0.032 (standard deviation 0.19). This low variance establishes a stable baseline against which test samples can be compared, supporting sensitive detection when tumor fractions are low.

**Figure 5.**
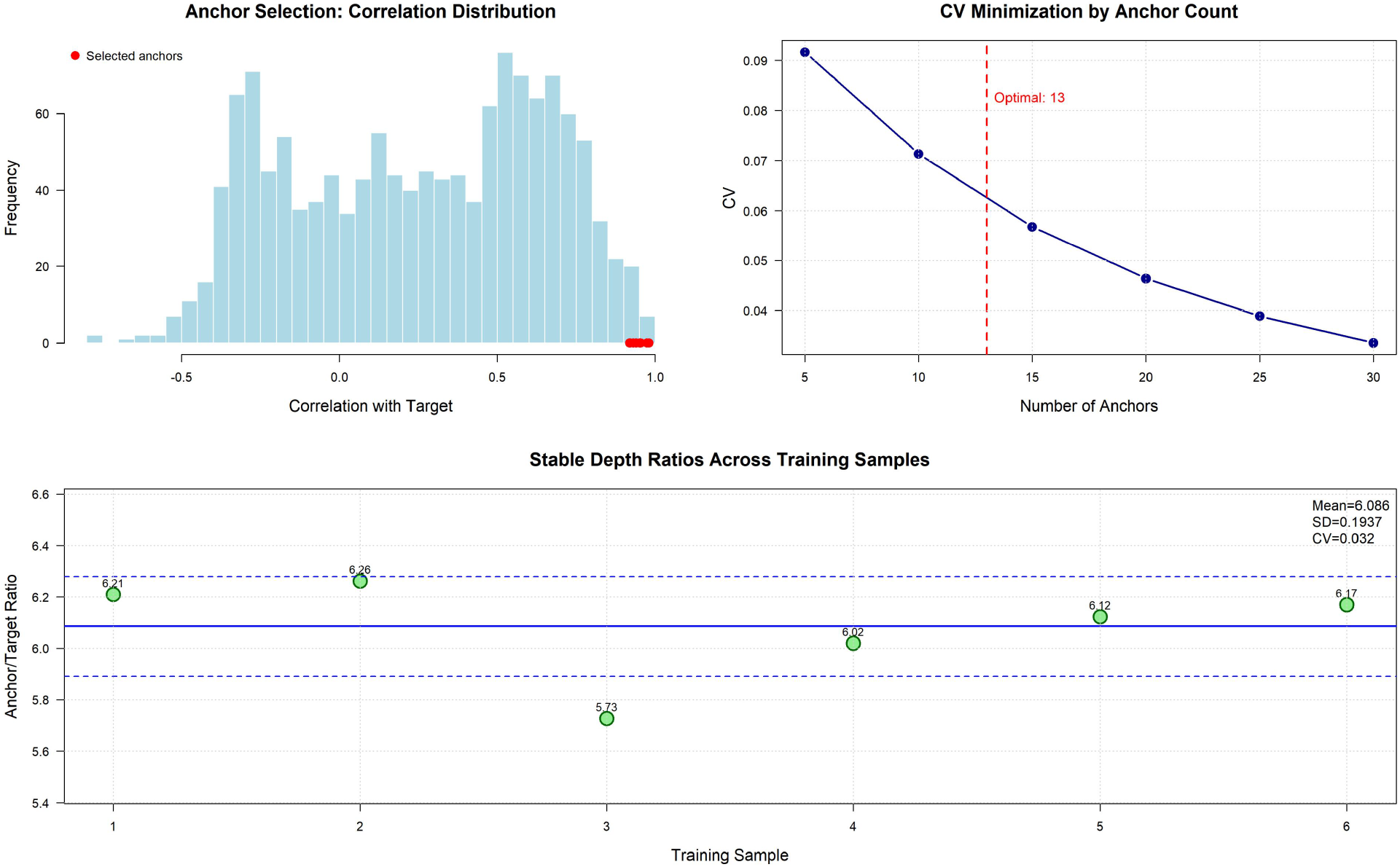
Anchor selection and recalibration process. (Insert Figure_A269_v5_anchor_selection.png here. Legend: Top-left: Pearson correlation distribution between a representative MET target segment and all other segments; red points indicate the selected anchors with the highest correlations. Top-right: coefficient of variation (CV) as a function of anchor count; the algorithm selects 13 anchors that minimize the CV. Bottom: anchor-recalibrated depth ratios across the six training samples, showing tight clustering (CV=0.032) that forms a stable baseline for CNV calling.)

### 3.5. Clinical Validation on a Real ctDNA NSCLC Cohort

To evaluate CESAR on real-world clinical samples, we applied the full four-step pipeline to a cohort of nine ctDNA samples from NSCLC patients, profiled on a 94-gene targeted panel. Based on prior clinical annotation, three samples carried MET amplifications, while the remaining six served as a matched panel of normals for training. On this cohort, the standard CNVkit-based pipeline did not distinguish the MET-positive samples from the negative baseline.

CESAR was trained on the six annotation-negative samples to establish baseline anchor sets and copy-number distributions. When applied to all nine samples, CESAR separated the three MET-positive cases from the six negatives (Figure 6). The three positive samples exhibited copy numbers of 1.82 to 3.30 with confidence scores of 13.9 to 100 (-log_10_ P), while the six negatives clustered around the diploid baseline (CN 1.95–2.06, confidence scores < 1.5). Notably, sample #3 (CN 1.82) showed only a modest copy-number increase yet was called with high statistical confidence (conf 32.5), illustrating CESAR’s sensitivity to subtle amplifications supported by consistent evidence across segments. This separation shows that CESAR recovers clinically annotated MET amplifications in real ctDNA data where the standard pipeline did not.

**Figure 6.**
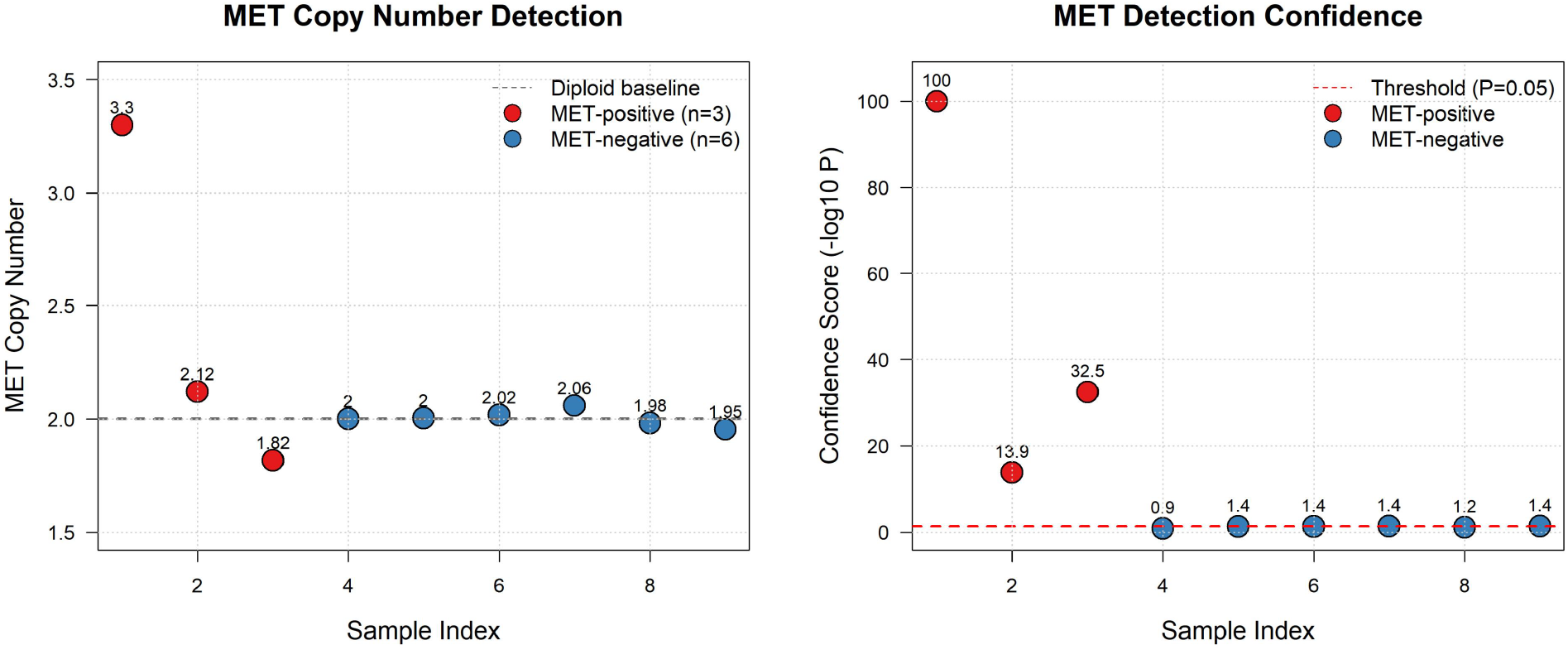
CESAR MET detection on a clinical ctDNA NSCLC cohort. (Insert Figure_A269_MET_FINAL.png here. Legend: left panel, MET copy number for nine NSCLC ctDNA samples; right panel, corresponding confidence scores. Red points indicate the three annotation-positive samples; blue points indicate the six matched negatives. CESAR separates all three MET-positive samples from the diploid baseline, including sample #3, which shows a subtle CN increase (1.82) but strong statistical evidence (conf 32.5).)

To examine CESAR’s segment-level resolution, Figure 7 displays the copy-number calls for all 19 segments within the MET gene across the nine samples. The three positive samples consistently show elevated copy numbers across nearly all segments, while the six negatives remain at the diploid baseline. This segment-by-segment concordance indicates that the MET signal is not driven by individual outlier probes but reflects a gene-wide alteration. It also illustrates why the segment-based approach is robust to probe-level noise: even if a few segments give spurious values, the gene-level call integrates evidence across all segments.

**Figure 7.**
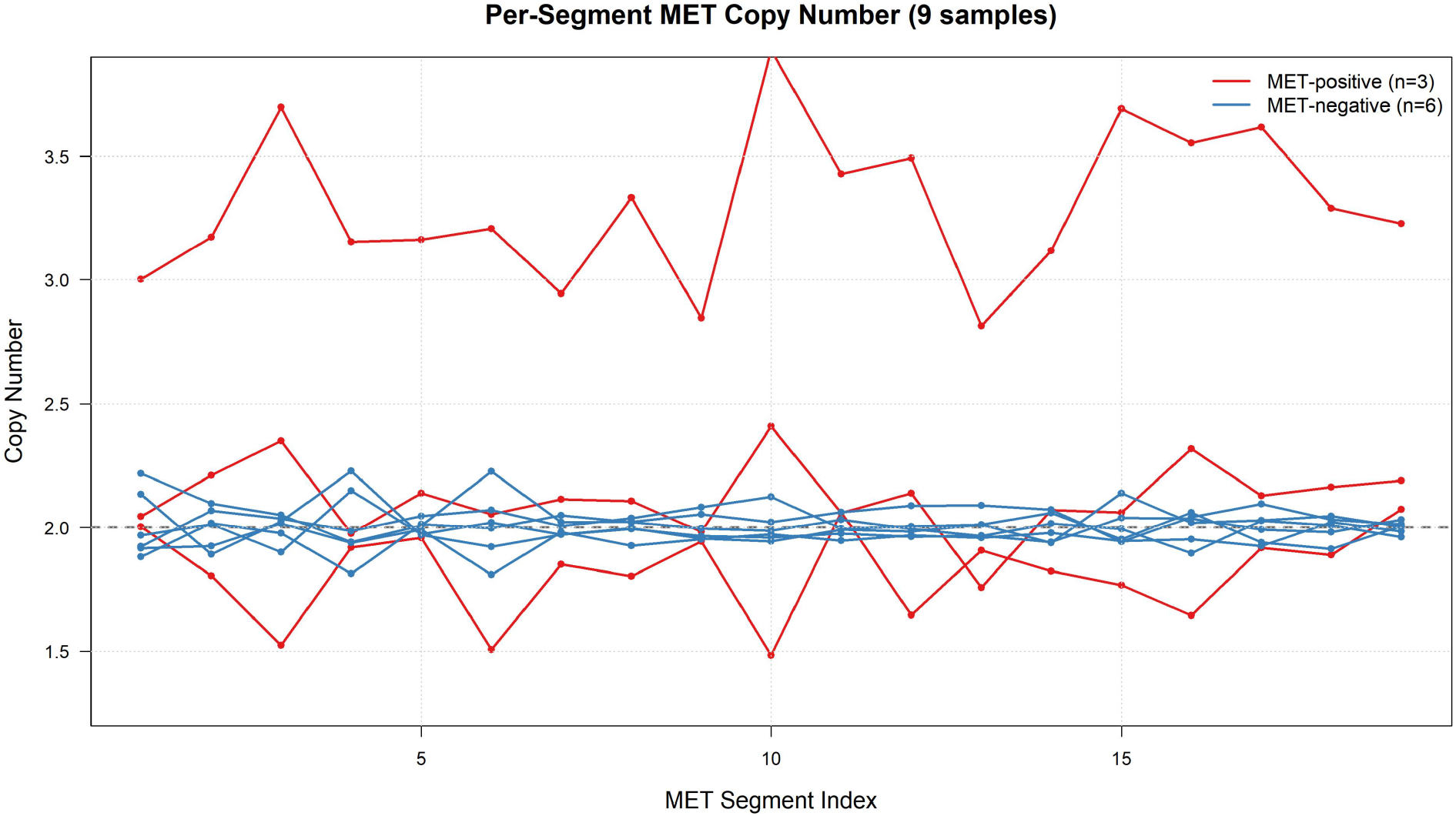
Per-segment MET copy number across nine samples. (Insert Figure_A269_v6_MET_segments.png here. Legend: copy-number estimates for all 19 MET segments. Each line represents one sample; red/orange lines indicate the three MET-positive samples, blue lines the six negatives. The segment-level concordance indicates a consistent, gene-wide amplification rather than an artifact of individual probes.)

This clinical validation is fully reproducible: the complete dataset (nine samples’ mpileup files), the segmentation BED file, trained model parameters, and a runnable end-to-end script are provided in the public GitHub repository (see Code Availability). The example can be re-executed from raw depth data through segmentation, anchor training, and final detection without external dependencies.

## 4. Discussion

Target enrichment sequencing relies on deploying numerous hybridization probes across the region of interest. Individual probes exhibit heterogeneity in capture sensitivity, binding stability, and variance driven by sequence context (e.g., GC content, secondary structure). As a result, CNV calling algorithms that rely on the global average depth of all probes within a target region are limited: their sensitivity is degraded by the inclusion of low-quality or highly variable probes.

The primary contribution of CESAR is to turn this probe-level heterogeneity to advantage. By employing the CBS algorithm, CESAR partitions the target space based on empirical capture behavior rather than arbitrary genomic coordinates. The anchor recalibration step then replaces the global average—which is degraded by low-quality and highly variable probes—with a set of stable, highly correlated anchor segments that represent the baseline expected depth. Consistent with prior work on targeted-capture biases [13], substituting the noisy global average with these dynamically selected anchors reduces systematic bias, and does so without the computational cost of sequence-context-based denoising methods.

The sensitivity achieved by CESAR—resolving amplifications as low as 2.18 copies (1.09x fold change) on reference standards—is relevant for liquid biopsy, where the tumor signal is highly diluted. While B-allele frequency (BAF) and breakpoint-based methods offer high theoretical sensitivity, their utility is limited in focused clinical panels (e.g., 30 kb footprints) that rarely capture enough heterozygous SNPs or fusion breakpoints. CESAR is designed to extract copy-number information from read depth alone, which is the signal most reliably available in such panels.

The clinical applicability of CESAR was validated on a nine-sample NSCLC ctDNA cohort profiled on a 94-gene panel. In this real-world dataset, CESAR successfully identified three MET-amplified samples with high confidence (CN: 1.82–3.30, confidence scores 13.9–100), clearly separating them from six matched negatives clustered at the diploid baseline (CN: 1.95– 2.06). Importantly, the standard CNVkit-based pipeline failed to distinguish these groups, highlighting CESAR’s advantage in detecting clinically actionable alterations in challenging ctDNA contexts. The fact that this validation cohort is fully reproducible—with all raw data, trained models, and analysis scripts publicly available—allows independent verification of CESAR’s performance.

### 4.1. Limitations

Several limitations qualify these findings. First, the reference-standard evaluation used small per-condition sample sizes (n = 6–10 per batch); while sufficient to demonstrate reproducibility relative to CNVkit, larger cohorts would be needed to establish detection limits and confidence intervals with precision. Second, the clinical validation cohort comprised only nine NSCLC samples; although this dataset successfully demonstrated CESAR’s ability to detect MET amplifications where CNVkit failed, validation on larger and more diverse clinical cohorts is needed to assess generalizability across tumor types, panels, and sequencing platforms. Third, CESAR reports copy number relative to a learned anchor baseline rather than an absolute measurement; a relative loss may reflect either a true deletion at the target or a coordinated shift in the anchor segments, so apparent deletions in particular warrant orthogonal confirmation by methods such as ddPCR or FISH. Fourth, the current statistical model assumes the recalibrated ratios are approximately Normal and reports per-gene confidence without a formal multiple-testing correction across genes; panels interrogating many loci may require an added correction. Finally, CESAR depends on a well-matched panel of normals: performance degrades when the training normals do not reflect the capture chemistry, sequencing platform, or biofluid of the test sample. Addressing these points—orthogonal validation, larger cohorts, diverse tumor types, and explicit absolute-copy-number calibration—is the focus of ongoing work.

---

## 5. Conclusion

CESAR is a sensitive and reproducible computational framework for gene-level CNV detection in targeted panel sequencing of ctDNA. By combining relative-efficiency segmentation with individualized anchor recalibration, it improves the signal-to-noise ratio of depth-based calling in liquid-biopsy data. On reference standards it detects low-level driver-gene amplifications (e.g., MET, ERBB2, EGFR) with lower bias and variance than CNVkit, and on a clinical NSCLC ctDNA cohort it successfully identified MET amplifications where the standard pipeline failed. The complete pipeline, including raw data and trained models from the clinical validation, is publicly available and fully reproducible. With validation on larger and more diverse cohorts, CESAR is a promising tool for identifying actionable copy-number alterations in precision oncology.

---

## Code Availability

CESAR is distributed as an R package, `CesaR`, implementing the four-step pipeline described here (`build_master_depth`, `cesar_segment`, `cesar_train`, `cesar_detect`). The complete source code, user documentation, a bundled example dataset (`cesar_demo`, a spike-in dilution series covering the reference-standard design of Table 1), and a runnable real-ctDNA example are freely available in the public GitHub repository: [https://github.com/nishuai/CesaR](https://github.com/nishuai/CesaR). The bundled dataset and example scripts reproduce the segmentation, anchor-selection, and detection steps end to end without external data.

